# POMC neurons modulate infant vocalizations through opioid signaling

**DOI:** 10.1101/2022.08.15.504046

**Authors:** Gabriela M. Bosque Ortiz, Marcelo O. Dietrich

**Affiliations:** Laboratory of Physiology of Behavior, Department of Comparative Medicine, Yale School of Medicine, New Haven, CT, USA; Interdepartmental Neuroscience Program, Biological and Biomedical Sciences Program, Graduate School in Arts and Sciences, Yale University, New Haven, CT; Yale Center for Molecular and Systems Metabolism, Yale School of Medicine, New Haven, CT; Department of Neuroscience, Yale School of Medicine, New Haven, CT, USA

## Abstract

Infants cry when separated from their mothers. Here, we show that POMC neurons in the arcuate nucleus of the hypothalamus—via ß-endorphin production and signaling—modulate the crying behavior of mouse pups. The effect of POMC neurons on vocal behavior depends on the expression of µ-opioid receptors, the main receptor for ß-endorphin. Thus, POMC neurons in the hypothalamus modulate infant cry through opioid signaling.

## Main text

Infant mammals physically and emotionally attach to their mother or caregivers ^1^. This attachment provides most of the infant needs to thrive. As a consequence of this attachment, when separated from their mothers, infants respond with distress and protest, which include the emission of vocalizations—the infant cry response (or separation-induced vocalizations).

Infant mice and rats emit separation-induced vocalizations in the ultrasonic range ^2^, which can be used as a simple measurement of the infant’s attachment to the dam ^3^. The emission of separation-induced ultrasonic vocalizations (USV) has communicatory value as it directs maternal behavior toward the vocalizing pup ^4-7^. Despite the importance of this behavior for the development of the infant and for the formation and maintenance of the attachment bond between the infant and the mother, the neurons in the brain and their afferent signaling critical in the emission of separation-induced vocalizations remain elusive.

Endogenous opioids appear to play a key role in attachment behavior and in separation-induced vocalizations. For example, the expression and function of µ-opioid receptors affect attachment behavior in mice ^8^, primates ^9^, and likely humans ^10^. In addition, the injection of morphine—an opioid receptor agonist—reduces, while the injection of naloxone—an opioid receptor antagonist— increases the emission of separation-induced vocalizations ^11,12^. The source of these endogenous opioids that act on the µ-opioid receptor to modulate infant attachment behavior and cry is unknown.

Neurons in the hypothalamus that produce endogenous opioids might be key to the modulation of vocal behavior. POMC neurons in the arcuate nucleus are the main source of ß-endorphin— the endogenous opioid with the highest affinity for the µ-opioid receptor ^13,14^—in the brain ^15-17^. We, therefore, put forward the hypothesis that POMC neurons in the hypothalamus modulate the emission of separation-induced vocalizations through opioid signaling.

To test our hypothesis, we first used a loss-of-function approach. We characterized the effects of the ablation of POMC neurons in the emission of USVs in ten-day-old mouse pups (P10) (**Fig 1a**). In newborn *Pomc*-Cre pups, we injected an adeno-associated virus (AAV) that expresses the diphtheria toxin subunit A only in cells that express *Cre* recombinase (AAV-mCherry-DIO-DTA). The expression of DTA on POMC neurons leads to significant cell ablation (**Fig 1b**).

**Figure 1.**
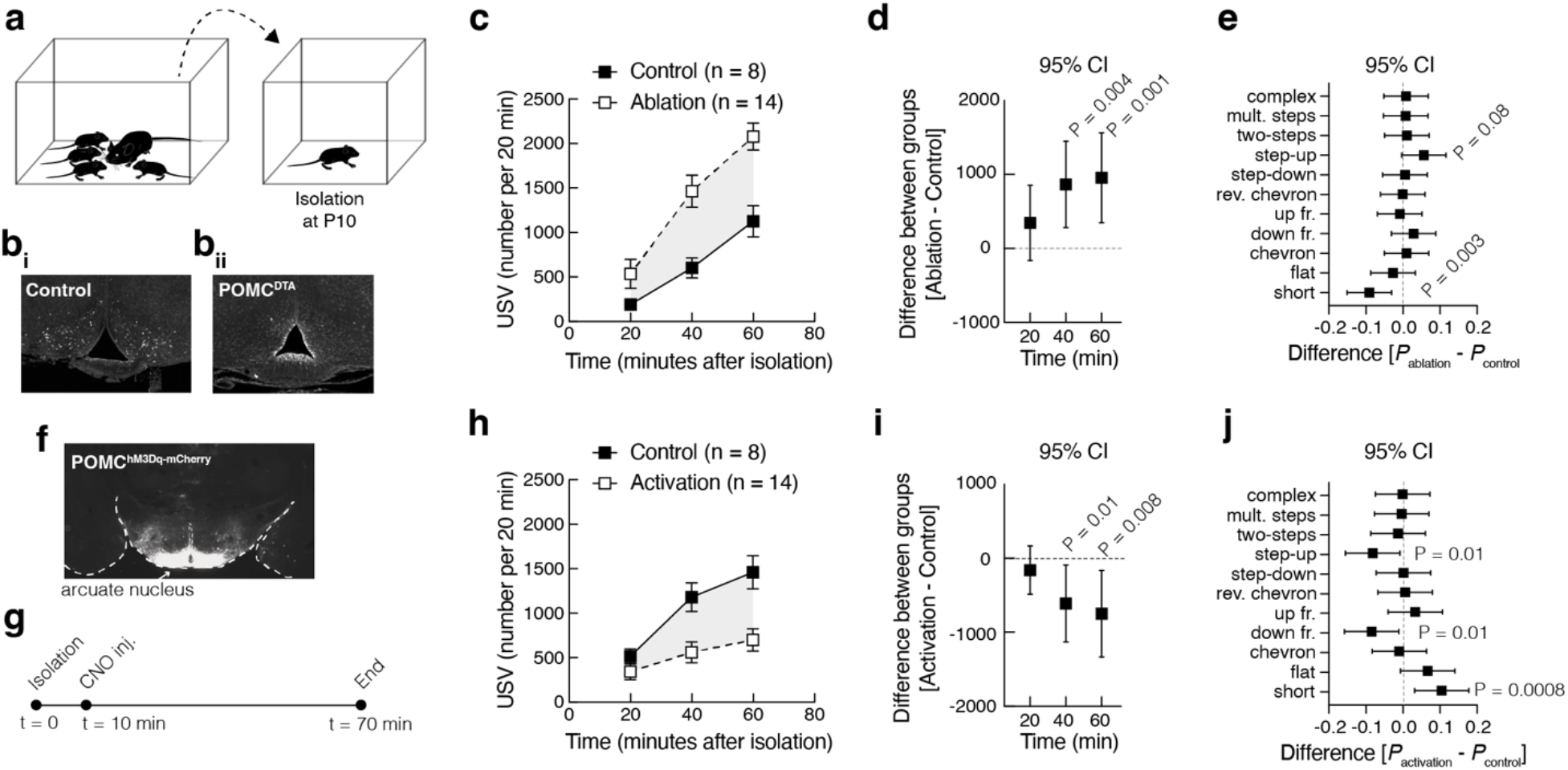
Ablation of POMC neurons increases, while activation decreases, the emission of separation-induced vocalizations in mouse pups. (**a**) Experimental design: ten-day-old mouse pups are isolated from the nest in a recording chamber; ultrasonic vocalizations are recorded and quantified. (**b**) Representative images of the arcuate nucleus of a control wild-type mouse (**bi**) and a *Pomc*-Cre mouse (**bii**), both injected with an AAV to express DTA in *Cre*-positive cells; expression of DTA ablates POMC neurons. (**c**) Number of USV emitted during isolation from the nest quantified in 20 min bins (symbols represent mean ± sem); statistical analysis using two-way ANOVA with time as a repeated measure and group as a factor (time: *F*_1.37, 27.43_ = 125.08, *P* < 10^−12^; group: *F*_1,20_ = 15.29, *P* = 0.008; interaction: *F*_2,40_ = 8.82, *P* = 0.0006). (**d**) Šídák’s multiple comparisons test; symbols represent 95% confidence intervals of the differences between groups. (**e**) Similar to (d) but showing the 95% confidence intervals of the difference in usage of vocal classes (normalized probabilities of all vocal classes sum to *P* = 1 per group); statistical analysis using two-way ANOVA with class and group as factors (class: *F*_10, 220_ = 120.4, *P* < 10^−15^; group: *F*_1, 220_ = 1.05, *P* = 0.99; interaction: *F*_10, 220_ = 3.03, *P* = 0.001). (**f**) Representative image of the arcuate nucleus of a *Pomc*-Cre mouse pup injected with an AAV-DIO-hM3Dq-mCherry at P0 (stained for mCherry). (**g**) Schematic of the protocol used to record the emission of USVs upon activation of POMC neurons: after 10 minutes of isolation, all pups are injected with CNO (1 mg/kg, i.p.) and recorded for an additional 60 minutes. (**h**) Similar to (c): statistical analysis using two-way ANOVA with time as a repeated measure and group as a factor (time: *F*_1.75, 35.16_ = 25.09, *P* < 10^−6^; group: *F*_1, 20_ = 7.54, *P* = 0.01; interaction: *F*_2, 40_ = 5.37, *P* = 0.008). (**i**) Similar to (d). (**j**) Similar to (e): statistical analysis using two-way ANOVA with class and group as factors (class: *F*_10, 220_ = 98.38, *P* < 10^−15^; group: *F*_1, 220_ < 10^−15^, *P* = 0.99; interaction: *F*_10, 220_ = 4.65, *P* < 10^−5^).

When P10, we separated pups from the next by placing them in a clean cage at room temperature and recording the emission of USVs for 60 minutes. Ablation of POMC neurons almost doubles the emission of USVs in P10 pups compared to controls (**Fig 1c-d;** statistical information is provided in the figure and figure legend). Using automated software to label each USV into one of eleven syllable classes ^18^, we revealed that ablation of POMC neurons leads to the emission of USVs of classes with more complex spectro-temporal features compared to non-ablated littermate control pups (**Fig 1e**).

To further interrogate the effects of POMC neurons on separation-induced vocalizations, we used a chemogenetic tool to artificially increase the activity of POMC neurons. We injected newborn *Pomc*-Cre pups with an AAV that expresses the excitatory receptor, hM3Dq, in cells expressing *Cre* (AAV-DIO-hM3Dq-mCherry) (**Fig 1f-g**). Wild-type littermate controls were also injected with the same virus but did not express the receptor. Injection of the synthetic ligand of hM3Dq, clozapine-N-oxide (CNO; 1 mg/kg, i.p.), activates POMC neurons in POMC^hM3Dq^ mice, but not in wild-type mice. Activation of POMC neurons in ten-day-old pups reduces by more than 50% the emission of USVs compared to control pups (**Fig 1h-i**) and leads to the emission of simpler classes of USVs (**Fig 1j**). Together, these results demonstrate that POMC neurons of the hypothalamus can bidirectionally modulate the emission of USVs in infant mice.

Because POMC neurons release ß-endorphin, we next tested the significance of ß-endorphin in the emission of separation-induced vocalizations. Towards this end, we used mice deficient in the production of ß-endorphin, but not for all other peptides derived from the cleavage of POMC. These mice harbor a point mutation in exon 3 of *Pomc*, which causes a premature stop codon and, consequently, a truncated POMC prohormone unable to produce ß-endorphin ^19^. Using these mice, we analyzed the emission of USVs from ß-endorphin null and wild-type pups within the same litter. Similar to the ablation of POMC neurons, pups deficient for ß-endorphin show higher rates of USVs compared to wild-type littermates (**Fig 2a-b**) and lower emission of USVs of the class short—the simplest form of USV (**Fig 2c**).

**Figure 2.**
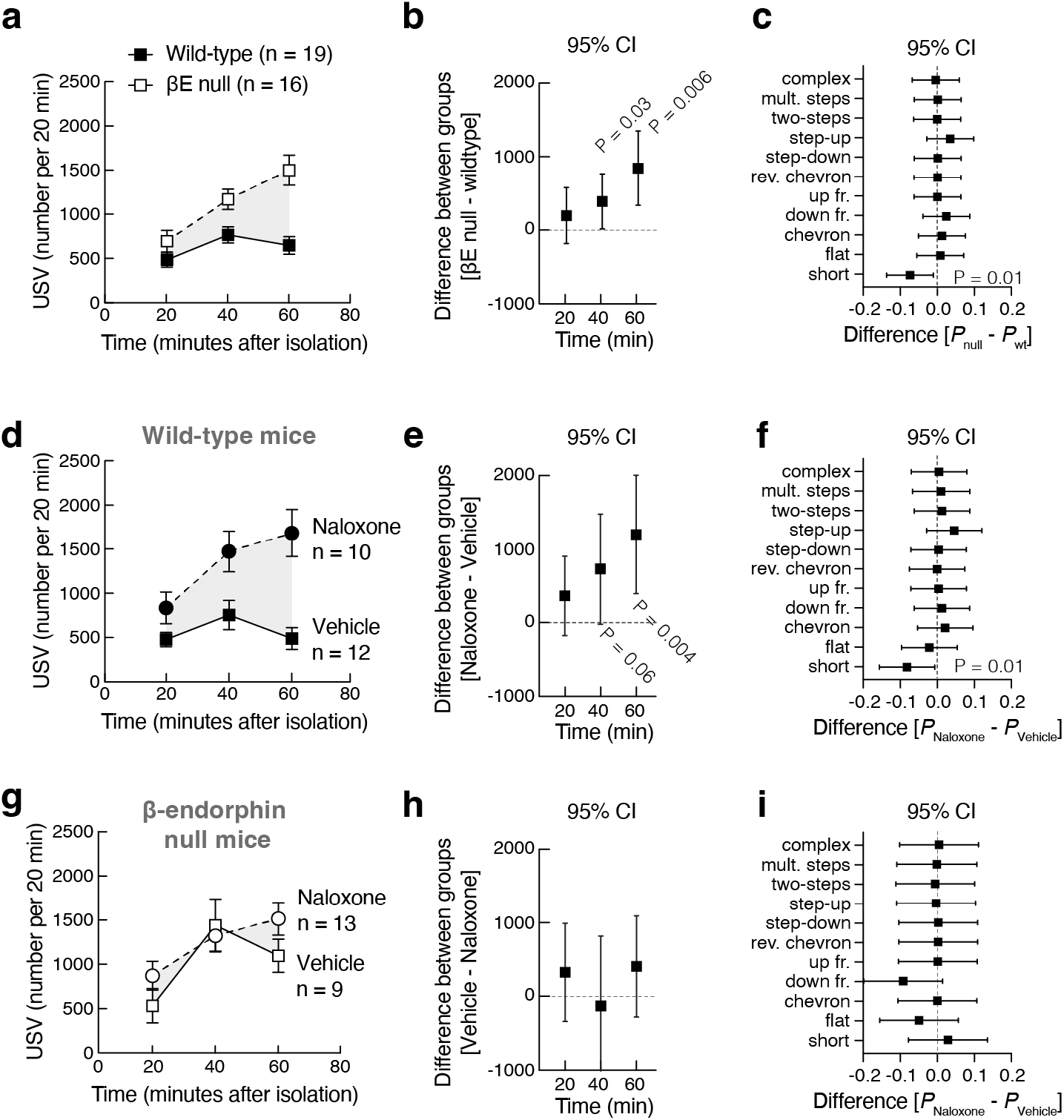
Loss-of-function of ß-endorphin increases separation-induced vocalizations in mouse pups. (**a**) Number of USV emitted during isolation from the nest in wild-type and ß-endorphin deficient mice quantified in 20 min bins (symbols represent mean ± sem); statistical analysis using two-way ANOVA with time as a repeated measure and group as a factor (time: *F*_1.22, 40.50_ = 18.10, *P* < 10^−4^; group: *F*_1, 33_ = 14.34, *P* = 0.0006; interaction: *F*_2, 66_ = 7.63, *P* = 0.001). (**b**) Šídák’s multiple comparisons test; symbols represent 95% confidence intervals of the differences between groups. (**c**) Similar to (b) but showing the 95% confidence intervals of the difference in usage of vocal classes (normalized probabilities of all vocal classes sum to *P* = 1 per group); statistical analysis using two-way ANOVA with class and group as factors (class: *F*_10,363_ = 138.13, *P* < 10^−15^; group: *F*_1, 363_ = 0.01, *P* = 0.88; interaction: *F*_10, 363_ = 1.42, *P* = 0.12). (**d**) Similar to (**a**), but data refers to wildtype mice treated with vehicle (saline solution) or naloxone (5 mg/kg, i.p.): statistical analysis using two-way ANOVA with time as a repeated measure and group as a factor (time: *F*_1.68, 33.61_ = 13.71, *P* < 10^−4^; group: *F*_1, 20_ = 11.45, *P* = 0.002; interaction: *F*_2, 40_ = 9.17, *P* = 0.0005). (**e**) Similar to (b). (**f**) Similar to (c): statistical analysis using two-way ANOVA with class and group as factors (class: *F*_10, 219_ = 77.00, *P* < 10^−15^; group: *F*_1, 219_ = 0.0001, *P* = 0.99; interaction: *F*_10, 219_ = 1.48, *P* = 0.14). (**g**) Similar to (**d**), but data refers to ß-endorphin deficient mice treated with vehicle (saline solution) or naloxone (5 mg/kg, i.p.): statistical analysis using two-way ANOVA with time as a repeated measure and group as a factor (time: *F*_1.74, 34.81_ = 21.20, *P* < 10^−5^; group: *F*_1, 20_ = 0.71, *P* = 0.40; interaction: *F*_2, 40_ = 3.20, *P* = 0.05). (**e**) Similar to (b). (**f**) Similar to (c): statistical analysis using two-way ANOVA with class and group as factors (class: *F*_10, 219_ = 48.52, *P* < 10^−15^; group: *F*_1, 219_ = 0.98, *P* = 0.32; interaction: *F*_10, 219_ = 0.77, *P* = 0.65).

To test whether the lack of signaling via opioid receptors plays a part in the altered vocal behavior observed in mice deficient for ß-endorphin, we treated wild-type and ß-endorphin null mice with naloxone, a non-selective and competitive opioid receptor antagonist that has the highest affinity for the µ-opioid receptor. The treatment of wild-type pups with naloxone increases the emission of USVs compared to saline-injected controls (**Fig 2d-e**) and lowers the emission of USVs of the class short—similar to the phenotype of ß-endorphin null mice (**Fig 2a-c**). In contrast, treatment of ß-endorphin null mice with naloxone did not significantly affect the emission of USVs (**Fig 2g-h**) or the type of USVs emitted (**Fig 2i**). Together, these results indicate that ß-endorphin, which is released by POMC neurons in the brain, modulates the emission of USVs in mouse pups via downstream opioid receptors.

Next, we tested the extent to which the effect of POMC neurons on the emission of separation-induced vocalizations depends on µ-opioid receptor signaling. We crossed *Pomc*-Cre mice with mice knockout for the µ-opioid receptor (*Orpm1*^*-/-*^). Within the same litter, we generated animals that express (or do not) *Cre* recombinase in POMC neurons and that were wild-type (*Orpm1*^+/+^) or knockout (*Orpm1*^*-/-*^) for µ-opioid receptor. Similar to the experiment above (**Fig 1**), we injected newborn mice with the AAV-DIO-hM3Dq-mCherry to express hM3Dq selectively in POMC neurons and to allow the artificial activation of these cells upon injection of CNO. In line with our previous experiment (**Fig 1**), activation of POMC neurons in *Orpm1* wild-type pups leads to a reduction in the emission of USVs (**Fig 3a-b**) and an altered vocal repertoire (**Fig 3c**). In contrast, activation of POMC neurons in *Orpm1*^−/-^ pups does not significantly change the emission of USVs (**Fig 3d-e**). Activation of POMC neurons in *Orpm1*^−/-^ pups, however, lowers the relative emission of USVs of the class short, instead of increasing as it occurs in wild-type pups (**Fig 1j** and **Fig 3c**). Thus, the expression of the µ-opioid receptor (*Orpm1*) is necessary for the quantitative and qualitative changes that occur in the emission of USVs upon activation of POMC neurons of infant mice.

**Figure 3.**
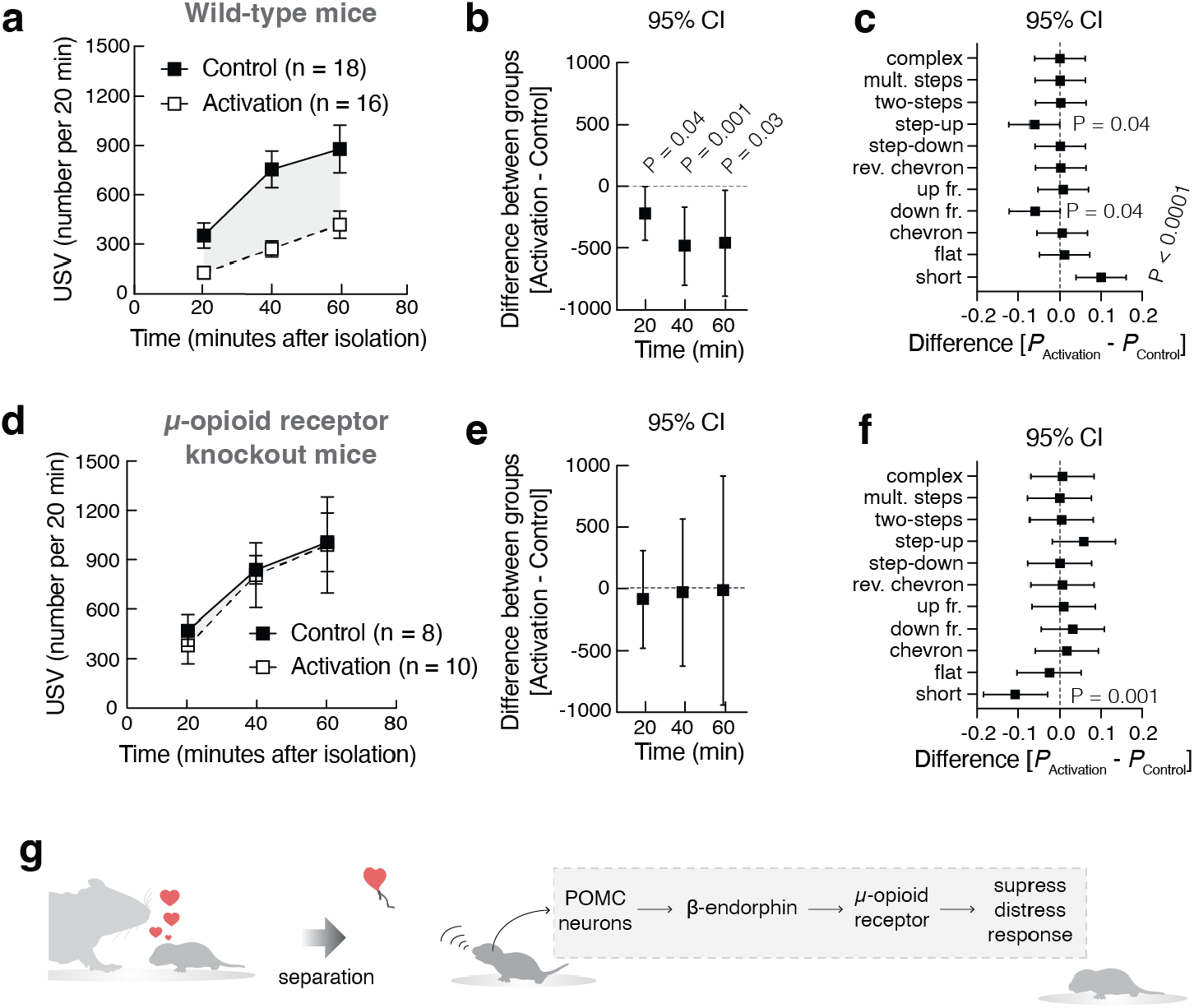
The expression of the µ-opioid receptor (*Orpm1*) is necessary for the suppression of USVs upon activation of POMC neurons. *Pomc*-Cre mice were backcrossed to *Orpm1* knockout mice and newborns were injected with an AAV-DIO-hM3Dq-mCherry. Wildtype and *Orpm1* knockout littermates were used for this experiment. At P10, after 10 minutes of isolation, all pups are injected with CNO (1 mg/kg, i.p.) and recorded for an additional 60 minutes for the emission of USVs. (**a**) Number of USV emitted during isolation from the nest in wild-type mice quantified in 20 min bins (symbols represent mean ± sem); statistical analysis using two-way ANOVA with time as a repeated measure and group as a factor (time: *F*_1.29, 41.31_ = 16.05, *P* < 10^−4^; group: *F*_1, 32_ = 13.85, *P* = 0.0007; interaction: *F*_2, 64_ = 1.97, *P* = 0.14). (**b**) Šídák’s multiple comparisons test; symbols represent 95% confidence intervals of the differences between groups. (**c**) Similar to (b) but showing the 95% confidence intervals of the difference in usage of vocal classes (normalized probabilities of all vocal classes sum to *P* = 1 per group); statistical analysis using two-way ANOVA with class and group as factors (class: *F*_10, 352_ = 162.26, *P* < 10^−15^; group: *F*_1, 352_ = 3.17, *P* > 0.99; interaction: *F*_10, 352_ = 3.82, *P* < 10^−5^). (**d**) Similar to (**a**), but data refers to *Orpm1* knockout mice: statistical analysis using two-way ANOVA with time as a repeated measure and group as a factor (time: *F*_1.10, 17.72_ = 8.66, *P* = 0.007; group: *F*_1, 16_ = 0.04, *P* = 0.82; interaction: *F*_2, 32_ = 0.03, *P* = 0.96). (**e**) Similar to (b). (**f**) Similar to (c): statistical analysis using two-way ANOVA with class and group as factors (class: *F*_10, 165_ = 92.21, *P* < 10^−15^; group: *F*_1, 165_ < 10^−15^, *P* > 0.99; interaction: *F*_10, 165_ = 2.35, *P* = 0.01). (**g**) Working model: during separation from the mother, POMC neurons of the hypothalamus release ß-endorphin that acts on downstream µ-opioid receptors to suppress the distress response of the infant and decrease the emission of separation-induced vocalizations.

Our results support a model in which—during separation from the mother—POMC neurons of the hypothalamus release ß-endorphin that acts on downstream µ-opioid receptors to modulate the emission of USVs (**Fig 3g**). Because the loss-of-function of POMC neurons and of ß-endorphin signaling increases the rate of vocalizations, our results suggest that the recruitment of POMC neurons/ß-endorphin signaling functions to suppress the distress response of the infant. This interpretation of our results finds support in previous studies investigating the effects of stress in adults. For example, the loss-of-function of ß-endorphin potentiates the behavior effects of stress ^20,21^. Moreover, stress activates POMC neurons ^22-24^ and the activation of POMC neurons is important for the behavioral and hormonal responses to stress ^22,23,25^. In aggregate, these findings support the role of ß-endorphin released by POMC neurons of the hypothalamus in the coping adaptations of the infant to the distress of separation.

Intertwined with POMC neurons in the arcuate nucleus are neurons that express agouti-related peptide (*Agrp*) ^26^. In the adult, ingestion of food activates POMC neurons, and the activation of POMC neurons promotes satiety. Conversely, ingestion of food inhibits Agrp neurons, and the activation of Agrp neurons promotes hunger. In infant mice, the activity of Agrp neurons during the separation of pups from the nest correlates with the modulatory role of these neurons in the emission of separation-induced vocalizations: activation of Agrp neurons increases, while the loss-of-function of these neurons decreases the number of vocalizations ^6^. Based on what we know about the function of these two populations of neurons in the adult brain, it is striking to realize that both POMC and Agrp neurons participate in the emotional responses of the infant to the mother ^27^, which appear to be independent of the provision of nutrition ^6,27^. Thus, at least in the mouse, the same neuronal populations that regulate food intake and energy balance in the adult, appear to be critically involved in the emotional attachment of the infant to their mother.

## Acknowledgments

We thank the members of the Dietrich laboratory and Tamas Horvath for their critical feedback on the project and the manuscript. We thank Jeremy Bober, Heidi Dong, and Shiwangi Goswami for their technical support. We thank David Gillich, Susan Andranovich, Valeria Krizsan, and Vickie Clark for administrative support. This work was only possible because of the availability of discretionary funds from the Yale School of Medicine that allowed the laboratory to take on new directions of research. The Dietrich laboratory received support from the National Institute of Mental Health of the National Institutes of Health under Award Number (R01MH125008), from the Foundation for Prader-Willi Research, and from the Odyssey Award from Smith Family Foundation. G.M.B.O. was supported by the HHMI Gilliam Fellowship. We thank David Bruin for copyediting the manuscript. The authors declare no conflict of interest.

## Author contributions

M.O.D and G.M.B.O. conceived the hypothesis, designed the study, and wrote the manuscript.

G.M.B.O. performed experiments and analyzed and plotted the data. The authors read and edited the manuscript.

## Data availability

Data (including raw data and audio files) are available on request from the corresponding author.

## Online Methods

### Experimental models and subjects

All preweaning mice used in the experiments were 9 to 10 days old from both sexes. Litters for POMC neuron activation and ablation were generated by breeding *Pomc*-Cre males (Jax #005965) with C57BL/6J (Jax #000664) females. Litters for POMC neuron activation in an *Orpm1* knockout background were generated by first breeding *Pomc*-Cre males with *Orpm1* knockout females (Jax #007559). *Orpm1* heterozygous males that were Cre positive were bred with heterozygous *Orpm1* heterozygous females. Litters for β-endorphin deficient mice (Jax #003191) were generated by breeding heterozygous mutants. In all experiments, littermates were used. Dams were 2 to 6 months old. All mice were kept in a temperature- and humidity-controlled room, in a 12/12 hr light/dark cycle, with lights on from 7:00 AM–7:00 PM. Studies took place during the light cycle. Food (Teklad 2018S, Envigo) and water were provided ad libitum. All procedures were approved by IACUC (Yale University).

### Viral injections

For activation of POMC neurons, we used an AAV8-hSyn-DIO-hM3D(Gq)-mCherry (Addgene #44361); for neural ablation we used an AAV1-Flex-taCasp3-TEVP (Addgene #45580). Newborns were removed from the home cage and placed in an aluminum vessel surrounded by ice for 10 minutes to induce hypothermia anesthesia. They are then placed in a stereotaxic frame with ear bars designed for neonates. A stereotaxic manipulator is used to position the syringe. Viral vectors (AAVs) were injected bilaterally at a volume of 0.3 µl per side using the following coordinates from lambda: AP = +.98 ML, lateral = -0.3mm, DV = -4.1.

### Immunohistochemistry

Mice were deeply anesthetized and perfused with freshly prepared fixative (paraformaldehyde 4%, in PBS 1x [pH = 7.4]). Brains were post-fixed overnight in fixative. Brain was sectioned at 100 µm in coronal orientation, allowing for efficient visualization of fibers extending through the brain without overlap. These sections were washed several times in PBS 1x (pH = 7.4) and pre-incubated with Triton X-100 (0.3% in PBS 1x) for 30 min. Sections were then incubated in a blocking solution (Triton 0.3%, Donkey Serum 10%, Glycine 0.3M in PBS 1x) for one hour. To confirm virus expression, brains were incubated with rabbit polyclonal anti-mcherry (1:1000; sc-52, Santa Cruz Biotechnologies) or chicken monoclonal anti-EGFP (1:1000, ab13970, abcam) for 16 hrs. After, sections were washed in 0.3% Triton in PBS, incubated with secondary fluorescent Alexa antibodies (1:500) for 4 hours, and washed once more with PBS. Sections were then mounted and imaged in a Keyence microscope or in a Leica SP5 confocal microscope.

### Protocol for separation-induced vocalization

Pups from the same litter were placed individually in a soundproof chamber containing fresh bedding material. An UltraSoundGate Condenser Microphone CM 16 (Avisoft Bioacoustics, Berlin, Germany) was placed 10 cm above the recording chamber and connected to the UltraSoundGate 416 USGH device to record ultrasonic vocalizations. Four to eight chambers were recorded simultaneously. After testing, mice were placed back in their home cage with the dam.

In the experiments with activation of POMC neurons, pups were recorded in isolation for 10 minutes (baseline) before receiving an intraperitoneal injection with hM3Dq ligand, clozapine-N-oxide (CNO, 1 mg/kg, dissolved in saline). Once injected, pups were placed back in the recording chamber for another hour. For naloxone experiments, pups were recorded for a 30-minute baseline and then injected with naloxone (5 mg/kg in saline, i.p) or saline (control); after, pups were placed back for another hour.

### Vocalization analysis

USVs were automatically extracted from the audio recordings by custom-built software using spectral analysis through image processing. The spectrograms were converted to grayscale images and the vocalizations were segmented on the spectrogram through a sequence of image processing techniques. The segmented vocalization candidates were then analyzed by a local median filtering to eliminate segmentation noise based on the contrast between a vocalization candidate and its background. Next, all the vocalizations were classified into 11 distinct call types by a Convolutional Neural Network, which had the AlexNet architecture as a starting point. The network was trained for USV classification with over 14,000 samples of real vocalizations, which were then augmented in order to increase the variability of the samples, resulting in >57,000 samples. Each vocalization received a label based on the most likely call type label attributed by the Convolutional Neural Network.

### Quantification and statistical analysis

Prism 8.0 or above was used to analyze data and plot figures. Shapiro-Wilk normality test was used to assess the normal distribution of the data. The data were analyzed using two-way ANOVA or mixed-effects analysis. Sidak’s multiple comparisons test was used to find post hoc differences among groups and to calculate the 95% confidence intervals to report effect size. In the text, values are provided as mean ± SEM. *P* < 0.05 was considered statistically significant and, when necessary and as described above, was corrected using Bonferroni’s method. Statistical data are provided in the text and in the figures.

